# Signal-space projection suppresses the tACS artifact in EEG recordings

**DOI:** 10.1101/823153

**Authors:** Johannes Vosskuhl, Tuomas P. Mutanen, Toralf Neuling, Risto J. Ilmoniemi, Christoph S. Herrmann

**Author notes:** Correspondence: Prof. Dr. Christoph Herrmann, Experimental Psychology Lab, Carl von Ossietzky Universität Oldenburg, Ammerländer Heerstr. 114-118, 26131 Oldenburg. These authors contributed equally to the article (Johannes Vosskuhl, Tuomas P. Mutanen).

## Abstract

1.

**Background:** To probe the functional role of brain oscillations, transcranial alternating current stimulation (tACS) has proven to be a useful neuroscientific tool. Because of the huge tACS-caused artifact in electroencephalography (EEG) signals, tACS–EEG studies have been mostly limited to compare brain activity between recordings before and after concurrent tACS. Critically, attempts to suppress the artifact in the data cannot assure that the entire artifact is removed while brain activity is preserved. The current study aims to evaluate the feasibility of specific artifact correction techniques to clean tACS-contaminated EEG data.

**New Method:** In the first experiment, we used a phantom head to have full control over the signal to be analyzed. Driving pre-recorded human brain-oscillation signals through a dipolar current source within the phantom, we simultaneously applied tACS and compared the performance of different artifact-correction techniques: sine subtraction, template subtraction, and signal-space projection (SSP). In the second experiment, we combined tACS and EEG on a human subject to validate the best-performing data-correction approach.

**Results:** The tACS artifact was highly attenuated by SSP in the phantom and the human EEG; thus, we were able to recover the amplitude and phase of the oscillatory activity. In the human experiment, event-related desynchronization could be restored after correcting the artifact.

**Comparison with existing methods:** The best results were achieved with SSP, which outperformed sine subtraction and template subtraction.

**Conclusions:** Our results demonstrate the feasibility of SSP by applying it to human tACS–EEG data.

## 2. Introduction

The goal of transcranial alternating current stimulation (tACS) is often the modulation of oscillatory brain activity and the concurrent demonstration of behavioral consequences of the intervention (Herrmann et al., 2013). Thus far, most studies combining tACS with electroencephalography (EEG) have demonstrated effects on oscillatory brain activity only by comparing the EEG before and after tACS (Zaehle et al. 2010; Vossen et al. 2015; Kasten and Herrmann 2017), because EEG data recorded during stimulation is contaminated by the huge tACS-generated artifact. As a workaround, behavioral effects found during application of tACS have been interpreted as a proxy of changes of brain oscillations (Neuling et al. 2012; Polanía et al. 2012; Cecere et al. 2015).

Correction of the tACS artifact in EEG recordings is more challenging as it is the case for magnetoencephalography (MEG). Due to the high spatial sampling, MEG studies on concurrent tACS online effects rely on the application of spatial filtering (a.k.a. beamforming) to deal with the artifact (Neuling et al. 2015; Kasten et al. 2018; Herring et al. 2019). These spatial filters achieve a strong, yet imperfect attenuation of the tACS induced electromagnetic that required additional correction, e.g. by contrasting two conditions with similar extent of the residual artifacts (Kasten et al. 2018; Herring et al. 2019). These residuals likely originate from non-linear modulations of the tACS artifact elicited by physiological processes in the human body (Noury et al. 2016; Noury and Siegel 2018), which also have to be taken into account in EEG recordings. The issue of tACS artifact correction in MEG data is discussed elsewhere (Neuling et al. 2015; Noury et al. 2016; Neuling et al. 2017; Noury and Siegel 2018; Kasten and Herrmann 2019) and will not be further addressed in this article.

Even though MEG might be better suited to analyze concurrent neurophysiology during tACS, EEG is a lot more common as a research method and thus it is desirable to have a method to suppress the artifact in EEG as well. Only a few studies so far have approached this issue (Helfrich et al. 2014; Voss et al. 2014; Dowsett and Herrmann 2016; Kohli and Casson 2019). While they represent milestones in tACS research, these studies also disclose a fundamental question: How can one assure that the brain responses of interest are not removed and that no residual artifact remains? To answer this question, it would be necessary to evaluate the performance of the artifact-correction procedure; however, this cannot be easily achieved when the brain activity to be recovered is virtually unknown. To tackle this issue, we conducted two experiments. First, we used a phantom to have full control over the “brain” signal and the “tACS” signal. Using pre-recorded human EEG as the source-current waveform in the phantom, we simultaneously applied tACS and compared different artifact-correction techniques. Second, the obtained results were used to demonstrate the feasibility of the artifact-correction performance in a human tACS–EEG experiment.

## 3. Experiment 1: Phantom study

### 3.1. Material and Methods

#### 3.1.1. Terminology

Although we used a phantom head in the first experiment, we will use terminology that has been established in human experiments in order to promote readability. For example, we will use “EEG” to refer to the recorded signal and “tACS” to refer to the application of sine-wave current to the phantom head’s outer layer (“scalp”).

#### 3.1.2. Experimental setup

The experimental setup is depicted in Fig. 1A. We used Matlab 2012b (The MathWorks Inc., Natick, MA, USA) on a laptop to control the delivery of pre-recorded EEG and the tACS signal to a digital-to-analog converter (USB-6229 BNC, National Instruments, Austin, USA). From here, the EEG signal was driven through a dipole source located inside a phantom head. The tACS signal was first sent to a battery-operated stimulator system (DC stimulator plus, Eldith, Neuroconn, Ilmenau, Germany) before being applied to the phantom head. The EEG that was recorded from the phantom head was stored for offline analysis.

**Figure 1.**
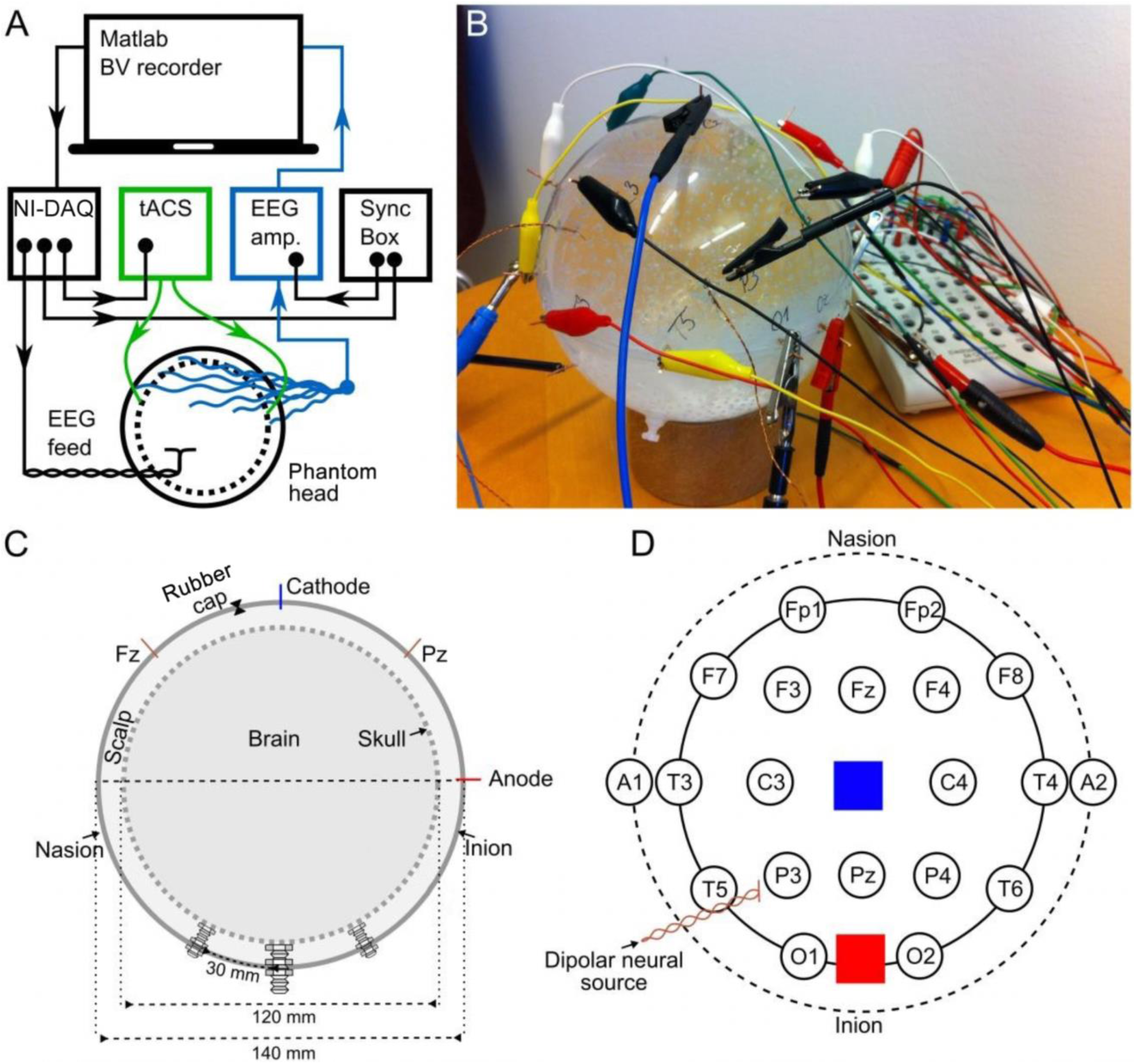
Experimental set-up and the structure of the phantom. A: Schematic illustration of the hardware setup as well as the signal delivery and recording (BV recorder: Brain Vision EEG recording software; NI-DAQ: National Instruments digital-to-analog converter). Arrowheads indicate the direction of information flow. B: Photo of the phantom head. C: Cross section of the phantom head along the midline. The dashed inner circle shows the inner porous spherical shell (skull), and the outer circle shows the outer spherical shell confining the scalp. Both the inner and outer shells were 2 mm thick. D: Stereographic projection of the phantom from above, depicting locations of EEG (circles) and tACS electrodes (blue and red squares), and the dipolar source.

#### 3.1.3. Phantom head

Our goal was to construct a phantom that captures crucial aspects of a human head receiving tACS: First, an artificial neural current source, second, a possibility to apply tACS to the surface of the phantom, and third, recording the combined signals. Furthermore, the phantom should possess the fundamental conductive properties of a human head: Most of the external current (tACS) is transmitted through the well-conducting skin, whereas the skull is a poor conductor of electricity. Likewise, most of the internal neuronally-driven ohmic currents remain inside the skull. Therefore, we built a spherical three-compartment phantom head with a dipolar current source inside the innermost space, as well as stimulation electrodes and recording electrodes on the outermost layer (Fig. 1B–D). The phantom head was filled with a fluid whose conductivity roughly matched that of human brain and scalp (0.57 S/m). The skull was realized as a porous spherical shell between the scalp and the brain with a conductivity of 0.019 S/m (conductivity values adapted from Gonçalves et al., 2003; Lai et al., 2005). A detailed description of the construction of the phantom head can be found in the supplemental information (S1.1).

#### 3.1.4. EEG

We delivered to the dipolar source of the phantom a signal waveform that resembles human EEG. Therefore, we used 60 s of human resting-state EEG previously recorded from a human participant (male, 24 years, right-handed) over the occipital cortex at electrode position O2 (Reference: nose) of the international 10–20 EEG system at 1 kHz sampling frequency using a Brain Vision Recorder (Brain Products, Munich, Germany). After up-sampling the signal to 100 kHz and high-pass filtering at 1 Hz, the signal was delivered to a dipole inside the phantom head and recorded from 18 electrodes (Fig. 1D) with 5 kHz sampling frequency amplified in the range of ± 3.2768 mV at a resolution of 0.1 μV (16 bits) using the Brain Vision Recorder with an online notch filter (50 Hz). The ground electrode was at location A1, the reference at a point comparable to the tip of the nose. The amplitude of the EEG signal driven through the dipole inside the phantom head was adjusted so that the amplitude of the resulting phantom EEG matched that of the pre-recorded human EEG (0.1–2.3 µV). To guarantee a perfect temporal alignment of the measured EEG at the phantom, the neural current source was synchronized with the tACS and the playback EEG via the BrainVision Syncbox (Brain Products, Munich, Germany; Fig. 1A). The EEG data were digitally stored for further offline analysis.

#### 3.1.5. tACS

We generated a digital 10-Hz sine wave at a temporal resolution of 100 kHz and output it via a digital-to-analog converter to the tACS electrodes of the phantom at electrode positions that were similar to Cz and Oz (Fig. 1D). The amplitude of the tACS signal was adjusted to avoid clipping, which would make the recovery of the EEG signal impossible. The largest tACS intensity that we could drive to the phantom without causing any clipping in the EEG channels was 150 µA, resulting in a maximum voltage between 30.5 and 1197.0 µV across the channels. We used two different tACS current intensities: 50 µA and 150 µA. The EEG amplitudes of the artifact depended linearly on the tACS current amplitude (50 µA: 10.2–399.3 µV) and were strongest in channels close to the tACS electrodes (Fig. 2B, left).

**Figure 2.**
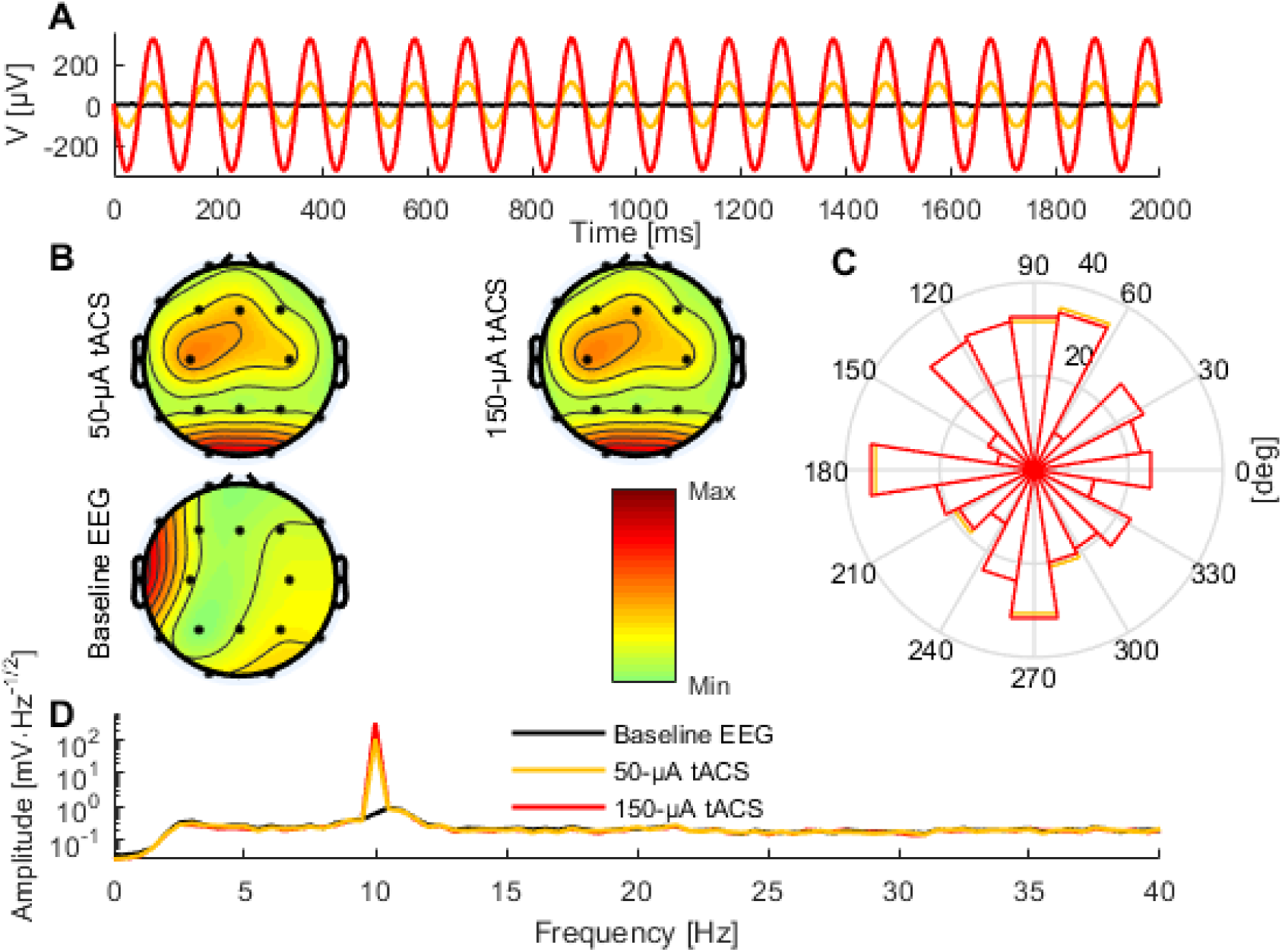
Comparison of baseline data and tACS-contaminated data. A: An illustrative 2-s segment of the baseline (black) and the artifactual data (red and yellow) measured at electrode Pz. B: The topographies showing the average amplitude in the range 8–12 Hz of the artifactual and baseline data. C: Circular histogram of the phase-difference distribution between the artifactual data and the baseline data at 10 Hz across all epochs and channels for the 50-µA (yellow) and 150-µA (red) condition. D: Frequency spectra of the 50-µA (yellow), 150-µA (red), and baseline conditions (black). Because of the logarithmic scale the peak amplitudes at 10 Hz for the 50-µA and 150-µA conditions appear similar regardless of the threefold difference.

### 3.2. Artifact correction

To evaluate the performance of different artifact-correction techniques, we first recorded the phantom head EEG resulting from the dipole current alone; this served as the baseline condition. Then, we applied tACS to the phantom while the dipolar source was active. Subsequently, we compared the performance of different artifact-rejection techniques (sine fitting, template subtraction, and signal-space projection (SSP)) in recovering the baseline signal from the data contaminated with the tACS artifact.

#### 3.2.1. Sine fitting

The most intuitive approach to remove the sinusoidal tACS-artifact is subtracting a sine wave at the stimulation frequency from the recorded data. This method has previously been applied to remove line noise from EEG data (Mitra & Bokil, 2007). We fitted a sine wave at the tACS frequency to non-overlapping time windows, each window having the length of one tACS period. The fitting was done for each channel separately, using the least-squares criterion with amplitude and phase as the fitted parameters. The resulting fits were then subtracted from the artifact-contaminated data in each time window.

#### 3.2.2. Template subtraction

The template subtraction method was adapted from a technique previously used to remove artifacts in simultaneous EEG / functional magnetic resonance imaging (fMRI) recordings (Allen et al., 2000; Niazy et al., 2005) and has also been applied to remove the tACS artifact from EEG data (Helfrich et al., 2014).

An artifact template was created by averaging data of a given number of tACS cycles and then subtracting the resulting template from each tACS cycle of the data. We used electrode-specific templates, which were obtained by averaging over all the tACS periods across the data segment of interest. These electrode-specific average artifact templates were then subtracted from the data in non-overlapping windows.

#### 3.2.3. Signal-space projection (SSP)

SSP is a method that separates signals into a set of different components that have constant spatial patterns in a multidimensional signal space, but whose amplitudes may change as a function of time; SSP has been used for separating, e.g., EEG and magnetoencephalography (MEG) signals (Uusitalo and Ilmoniemi, 1997). This feature can be exploited for the combination of EEG and tACS because tACS has a relatively constant spatial pattern, although it may change slightly due to changes in the conductive properties of the scalp. If we are able to estimate this spatial pattern accurately, we can use SSP to suppress the tACS artifact.

First, a maximally pure template of the artifact has to be calculated from the signal. To this end, single cycles of the sinusoidal tACS-artifact are averaged per channel. Thereby the brain signal is mostly removed from the recorded signal and only artifactual signals and noise remain. These remaining data are then used to estimate the artifact signal subspace, which then enables us to project out the artifact from the contaminated data. Here, the artifact subspace was estimated from the average artifact template (see section 3.2.2.), assuming that only little brain activity remains after averaging. The dimension of the artifact subspace was determined qualitatively from the singular value spectrum of the average artifact template. A detailed description of the SSP method can be found in the supplemental information (S1.2). One feature of SSP is that it introduces spatial distortions to the signal, which impedes conventional visual interpretation of the resulting signal. To minimize these undesired distortions and keep the corrected data visually interpretable, we used the source-informed reconstruction approach (SIR) introduced by Mutanen et al. (2016). The idea of SIR is to compute from the projected (distorted in a perfectly known manner) signal a brain current distribution and from this current distribution, the corrected signal. For SIR, one needs to compute the lead field matrix of the chosen forward model to explain the measured data in terms of source currents. We used a spherical model that had the same geometry as the phantom containing 5000 evenly distributed radial dipoles 50 mm away from the origin. From now on, we refer to the combined SSP–SIR approach simply as SSP. Since SIR is not sufficient to correct all the SSP-elicited distortions, we also applied SSP and SIR to the baseline data to make it comparable to the SSP cleaned data (further details in S1.2). The major benefit of this approach is that it allows a direct comparison of the two datasets (e.g. baseline data and artifact-contaminated data) because we are quantifying the change from the baseline to the tACS-contaminated data only in those signal-space dimensions that remain after cleaning. In essence, this approach takes into account the possible unwanted attenuation of the neuronal signals of interest (overcorrection).

#### 3.2.4. Analysis of the phantom data

To remove slow drifts and high-frequency noise, the data were bandpass-filtered from 2 to 80 Hz with a 4th-order Butterworth filter. For the artifactual datasets, we identified the exact data point when tACS started and the corresponding time point in the baseline dataset. We discarded the first second of data due to artifacts related to initializing the tACS and extracted a 50-s segment (1–51 s with respect to the tACS onset) for further analysis. We then applied the artifact-correction techniques to these data segments. The resulting data will be referred to as cleaned data. As indicated above, SSP was also applied to the baseline data prior to comparison. After the cleaned and the baseline data were divided into 2-s epochs, we Fourier-transformed each of these epochs and computed the epoch- and channel-specific amplitudes and phase-angle spectra.

To evaluate the performance of the artifact-rejection techniques, we estimated the degree of tACS-artifact contamination in the cleaned data. We calculated the residual artifact (*RA*) for each channel as:

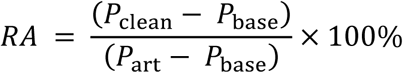

where *P*_clean_ *P*_base_, and *P*_art_ represent the signal power at 10 Hz for the cleaned, baseline, and artifactual data, respectively. A positive *RA* implies that the tACS artifact was not fully removed, whereas a negative *RA* suggests that some additional distortion was introduced in the data (e.g., attenuation of the signal of interest). We quantified spatial distortions elicited by the artifact rejection techniques by computing the topography maps of signal amplitude between 8 and 12 Hz of the baseline and cleaned data. We then computed the relative error (*RE*) between the baseline and the cleaned topographies:

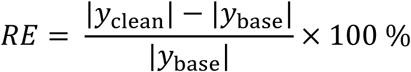

where *y*_clean_ and *y*_base_ are the topography vectors of the cleaned and the baseline data, respectively, and the |•| represents the Euclidian norm of the topography vector. The level of temporal distortions caused by the artifact-suppression methods was assessed by computing the correlation coefficient (CC) between the baseline and cleaned time courses in each channel and trial.

To further evaluate whether the amplitude spectrum of the neural source was recovered correctly, we computed the average spectrum over the epochs and compared the cleaned and baseline data of each channel separately. We focused the analyses on the individual alpha frequency (IAF: 10.5 Hz), the spectral peak in the range of 8–12 Hz estimated from the baseline spectrum. To test whether the correction methods distort the IAF amplitude, we performed a 2-way ANOVA (factor 1: 18 channels, factor 2: two conditions, i.e., cleaned data and baseline data) and post-hoc t-tests.

To analyze possible phase distortions, we subtracted for each epoch, channel, and frequency the baseline phase angle from the phase angle of the cleaned data. We visualized the phase difference at 10 Hz, when the artifact was in its maximum. Additionally, we computed the phase-locking value (PLV) (Lachaux et al., 1999) between the baseline and the cleaned data for each channel. To test whether the phase locking between the corrected and the baseline EEG was significant at the IAF, which would indicate preserved phase information, we used Bonferroni-corrected bootstrapping tests (Lachaux et al., 1999). All analyses were done using Matlab 2014b (The MathWorks Inc., Natick, MA, USA) and the EEGLAB 13.4.4b toolbox (Delorme & Makeig 2004).

### 3.3. Results

All artifact-correction methods were able to attenuate the tACS artifact (Table 1). After applying each method, the amplitude spectra were in the range of the signal of interest (Fig. 4); however, seemingly at the expense of different degrees of overcorrection, which means that also non-artifact activity had been removed. Sine fitting demonstrated higher overcorrection compared to template-subtraction and SSP (Table 1, RA results). The spatial information was best recovered by SSP, demonstrating only minor errors compared to template subtraction, whereas sine fitting yielded strong deviations (Table 1, RE results). Specifically, sine fitting shows the largest deviation in the frequency range of 5–15 Hz (Fig. 3, right). On average, SSP and template subtraction performed similarly; however, SSP had less variation across the channels, which can be seen in the more homogeneous topographies of SSP in comparison to Template subtraction (Fig. 3). Note the comparatively large *RE* in channel P3 after applying SSP, caused by a very low signal-to-noise ratio in this channel as can be already seen in the baseline condition (see Fig. 2B). Channel-wise frequency spectra further demonstrate the poor performance of the sine fitting within the 5–15-Hz range (Fig. 4, left). SSP yielded the best results, especially when comparing SSP-baseline data with SSP-cleaned data: the spectra matched almost perfectly (Fig. 4, right). Furthermore, SSP was the superior method in recovering the temporal information (phase) of the baseline signal, whereas template subtraction and sine fitting poorly recovered the baseline signal in a number of channels (Fig. 3, left). The difference in preserving the temporal information was also supported by high correlations of the signal between the baseline and the SSP-cleaned data compared to the other methods (Table 1, CC results). Additional support comes from the results of the bootstrapping analysis of the PLV at the IAF: After SSP, the PLV between cleaned and baseline data was significant, for all channels and conditions (*p*<0.05, after Bonferroni correction). After template subtraction, PLV was significant in most of the channels in both conditions (*p*<0.05) except for three cases (50 µV tACS - P3: *p*=1.62; 150 µV tACS - P3: *p*=14.58, O1: *p*=0.36, after Bonferroni correction). After sine fitting, no results were significant in the 50-µA-tACS condition and only one channel had a significant PLV in the 150-µA-tACS condition (F3: p<0.05, after Bonferroni correction).

**Table 1.**
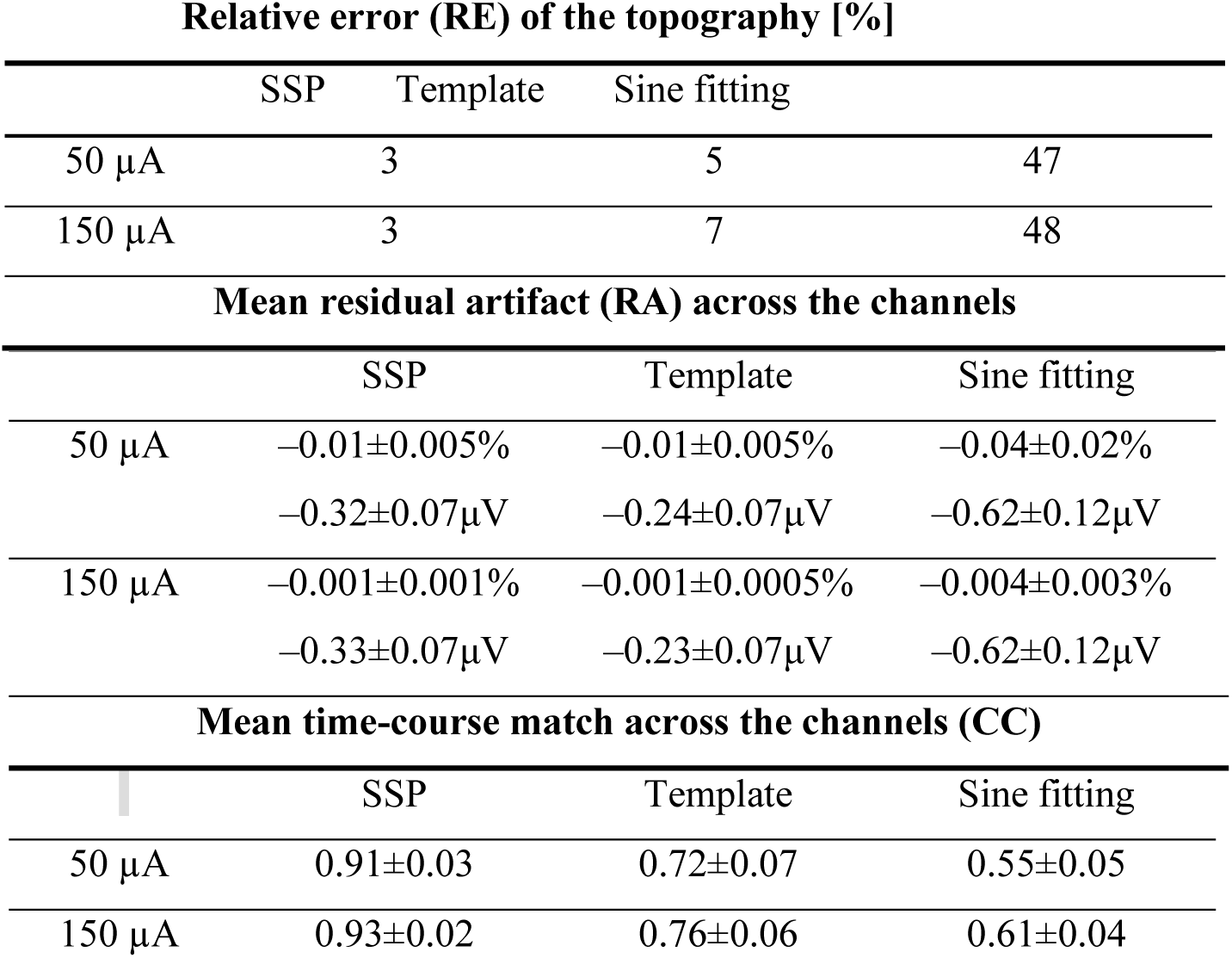
Comparison of the artifact-correction performance. Relative error (RE), Residual artifact (RA), and Correlation (CC). Note that a positive RA implies that the tACS artifact was not fully removed, whereas a negative RA indicates overcorrection.

**Figure 3.**
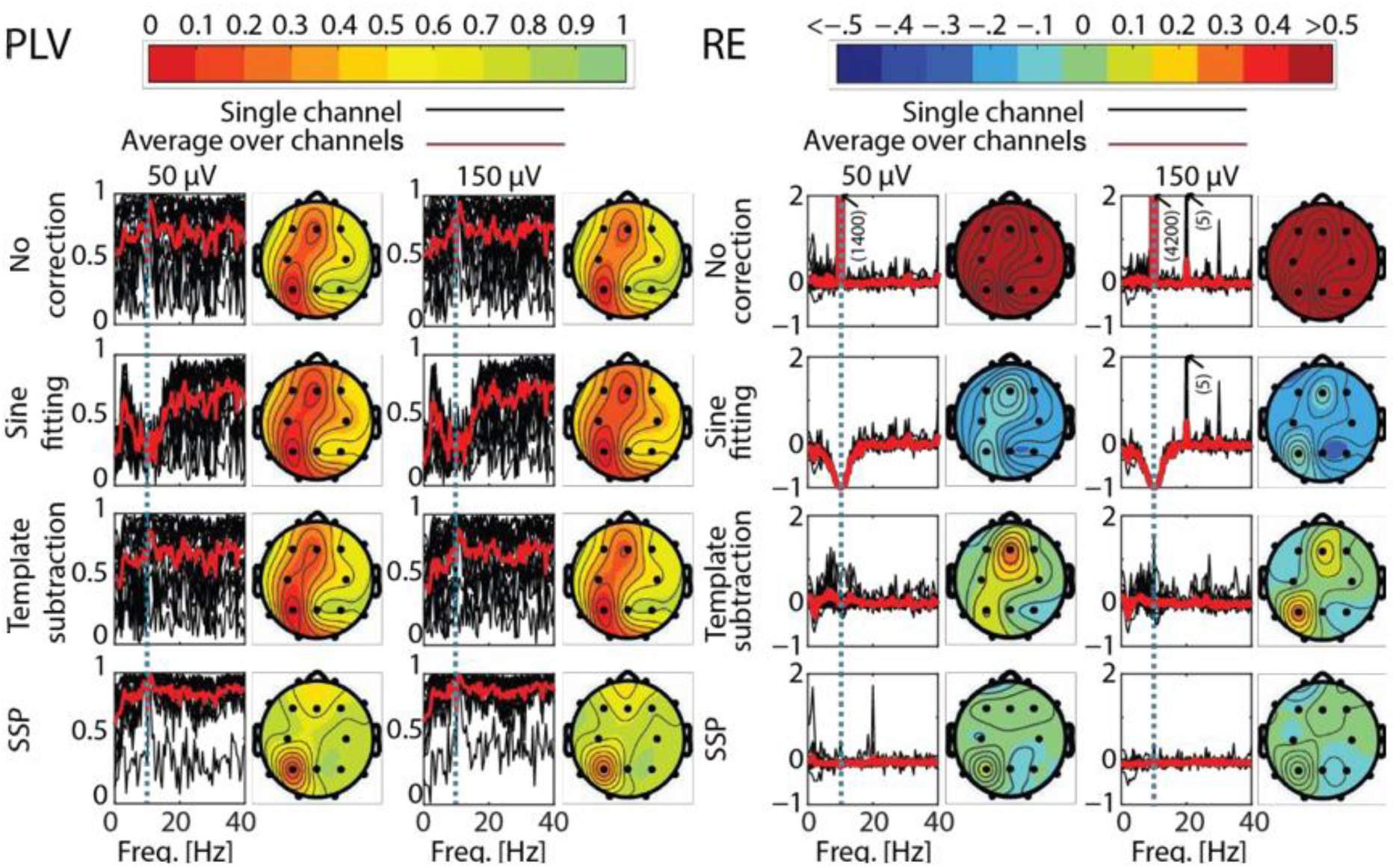
Phase-locking value (PLV) and relative error (RE) as function of frequency and different channels after artifact correction. PLV (left) and RE (right) for the different correction methods and tACS conditions as compared to the baseline. The black curves show PLV and RE as function of frequency in different channels, the red curve showing the average of the black curves. The dotted blue line depicts the stimulation frequency. Corresponding topographies show the mean value of the channel-specific black curves averaged across frequencies 0-40 Hz. In the PLV column, red indicates low PLV, meaning big distortions between baseline and corrected EEG. In the RE column, red means there was artifact left in the data, while blue depicts an overcorrection. Note that the color map of RE is thresholded to 50% absolute error and the plots in the RE section show only values between −1 and 2.

**Figure 4.**
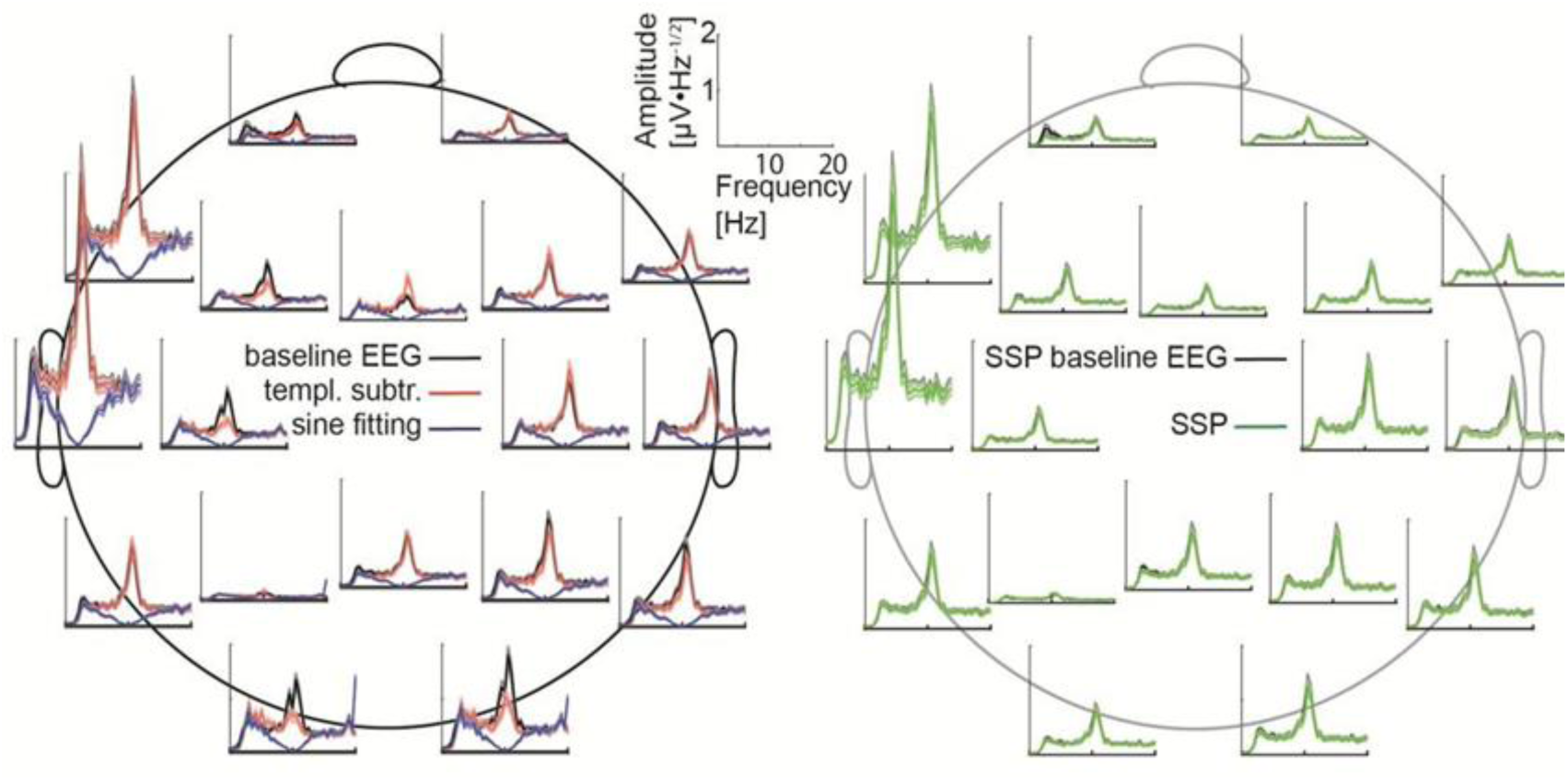
Frequency spectra for each channel. Left: The baseline data (black) are compared with the cleaned data after applying sine fitting (blue) and template subtraction (red). Right: SSP baseline data (black) compared with the SSP cleaned data (green). Here, 150-µV stimulation was delivered at 10 Hz.

For sine fitting, the 2-way ANOVA (factor 1: 18 channels, factor 2: two conditions, i.e., cleaned data and baseline data) demonstrated that the IAF amplitude depends on the tACS intensity, which additionally interacted with the channel (cf. Table 2). Furthermore, the interaction between the channel and condition was significant, which means the artifact suppression is not reliable. Subsequent post-hoc t-tests indicated significant changes in IAF amplitude in all channels and in both stimulation intensities (*t*(24)=8.96, *p*<0.001 for all channels).

**Table 2.** Results of the two-way ANOVA on the IAF amplitude changes. Asterisks mark statistically significant effects, i.e., the recovered signal deviates from the original signal.

|  | SSP | Template | Sine Fitting |
| --- | --- | --- | --- |
| **Main effect** | **50-μV:**<br>$F(1)=1.42$, $p=0.23$ | **50-μV:**<br>$F(1)=3.83$, $p=0.05$ | **50-μV:**<br>$F(1)=956$, $p<0.001^*$ |
| **Condition**<br>**(Baseline/Cleaned)** | **150-μV:**<br>$F(1)=1.21$, $p=0.27$ | **150-μV:**<br>$F(1)=4.43$, $p=0.04^*$ | **150-μV:**<br>$F(1)=959$, $p<0.001^*$ |
| **Main effect** | **50-μV**<br>$F(17)=78.29$, $p<0.001^*$ | **50-μV:**<br>$F(17)=77.24$, $p<0.001^*$ | **50-μV:**<br>$F(17)=40.78$, $p<0.001^*$ |
| **Channel** | **150-μV:**<br>$F(17)=78.74$, $p<0.001^*$ | **150-μV:**<br>$F(17)=76.53$, $p<0.001^*$ | **150-μV:**<br>$F(17)=40.76$, $p<0.001^*$ |
| **Interaction** | **50-μV:**<br>$F(1,17)=0.06$, $p=1$ | **50-μV:**<br>$F(1,17)=0.9$, $p=0.57$ | **50-μV:**<br>$F(1,17)=38.27$, $p<0.001^*$ |
| **Channel×Condition** | **150-μV:**<br>$F(1,17)=0.06$, $p=1$ | **150-μV:**<br>$F(1,17)=1.43$, $p=0.11$ | **150-μV:**<br>$F(1,17)=38.28$, $p<0.001^*$ |

Likewise, after template subtraction, the ANOVA revealed a significant main effect of condition in the 150-µA-tACS data, but no interaction between channels and conditions. No such main effect for condition was found in the 50-µA tACS condition. The ANOVAs on the SSP data neither showed effects for the condition nor for the interactions in both tACS conditions (Table 2). In general, SSP outperformed sine fitting and template subtraction. A summary of the SSP performance is depicted in Fig. 5.

**Figure 5.**
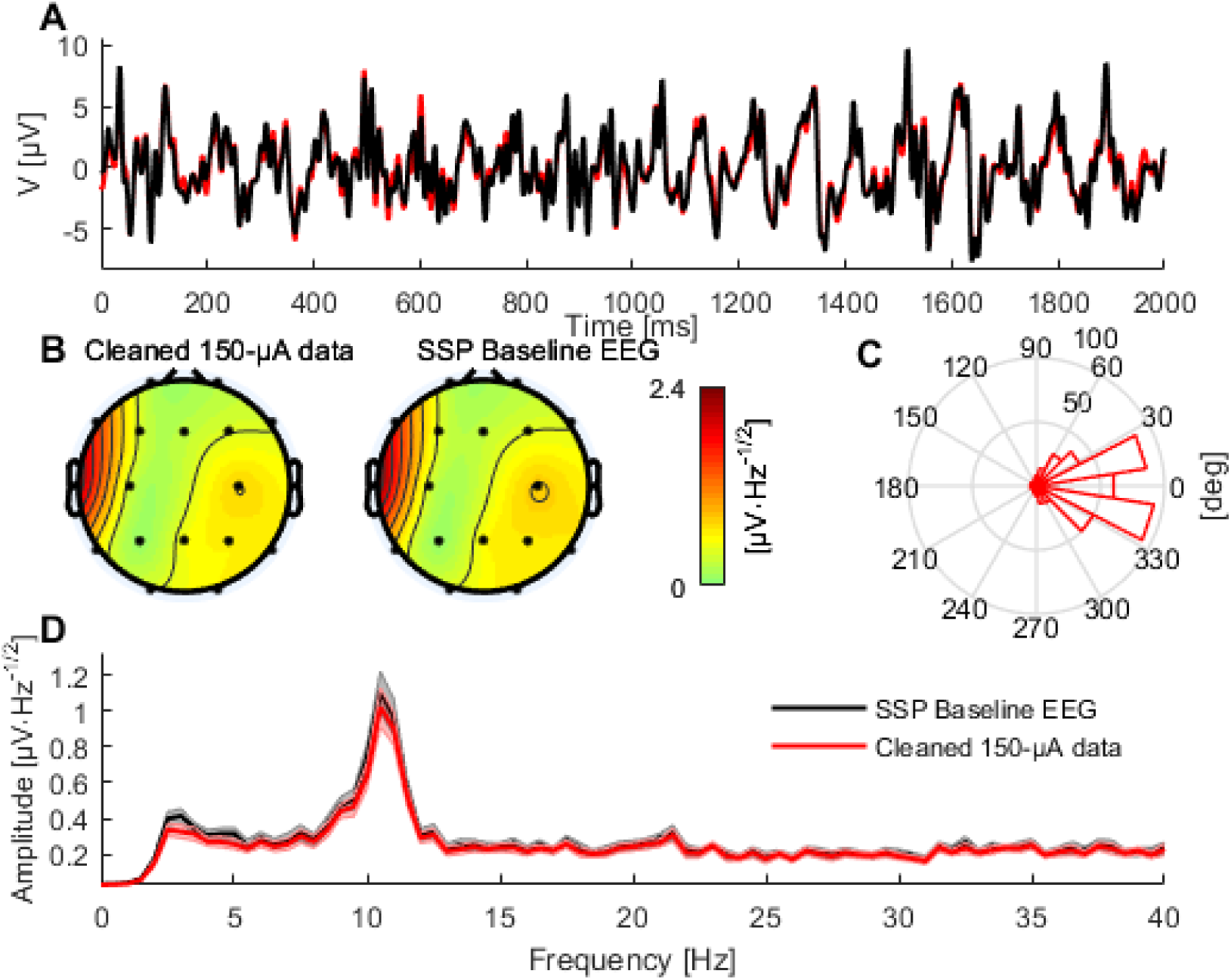
Comparison between baseline and SSP-cleaned data when delivering tACS with 150 µA. A: A 2-s segment of baseline (black) and cleaned data (red) measured at electrode Pz. B: Topographies showing the mean signal amplitude between 8 and 12 Hz of the cleaned and baseline data, respectively. C: Histogram of the phase difference between artifactual and baseline data at 10 Hz across all epochs and channels. D: The mean frequency spectra (averaged over epochs) of the baseline (black) and the cleaned data (red), in channel Pz. Shaded areas indicate the standard error of the mean.

## 4. Experiment 2: Human study

Demonstrating that a large portion of the tACS artifact can be suppressed with SSP in a phantom is only a first step towards correcting the tACS artifact in real human EEG data. To demonstrate that SSP is feasible for human tACS–EEG data, we recorded EEG-data during the application of tACS while the subject engaged in a mental rotation task. The mental rotation task (Shepard and Metzler 1971) is known to modulate ongoing alpha activity; while the stimuli are presented, occipital alpha oscillations desynchronize (Michel et al. 1994; Klimesch 1999). This event-related desynchronization (ERD; Pfurtscheller and Lopes da Silva 1999) has been used in studies to estimate the performance of methods for tACS artifact correction in MEG (Kasten et al. 2018). We thus employed a mental rotation task highly similar to Kasten and Herrmann (2017) and Kasten et al. (2018) to test the performance of the SSP-correction.

We applied tACS concurrently with a mental rotation task using an open-source stimulus set (Ganis and Kievit 2015, Fig. 6A). The performance of SSP to correct the tACS artifact can be assessed by its ability to recover the ERD expected during presentation of the visual stimuli. All experimental procedures were approved by a local ethics committee at the University of Oldenburg (Komission für Forschungsfolgenabschätzung und Ethik) and were in line with the Declaration of Helsinki. To achieve a high comparability with tACS intensities as used in many previous studies, we decided to apply tACS at 500 and 1000 µA. Note here that these tACS intensities might seem incomparable to the intensities used in the phantom (50 and 150 µA). The intensity of the stimulator output (in µA), however, is not as relevant when considering the correction of the artifact strength as measured via the EEG system (in µV). See table 3 for a comparison of these values for our study. The artifact strengths for the human study turn out to be higher by a factor of 20 compared to the phantom study. Thus the artifact correction in the human study is a lot more difficult.

**Table 3.** Comparison of the tACS intensities, as set in the stimulator (left column), and artifact strengths as measured in the EEG signal (right column) between the phantom and the human study. Artifact strengths differ considerable between channels. We thus report minimal and maximal values over the different channels in this table.

|  | Stimulator Output | Artifact strength<br>(min – max) |
| --- | --- | --- |
| **Phantom** | 50 $\mu\text{A}$ | 10 – 400 $\mu\text{V}$ |
| | 150 $\mu\text{V}$ | 30 – 1200 $\mu\text{V}$ |
| **Human** | 500 $\mu\text{A}$ | 30 – 10100 $\mu\text{V}$ |
| | 1000 $\mu\text{A}$ | 80 – 19600 $\mu\text{V}$ |

**Figure 6.**
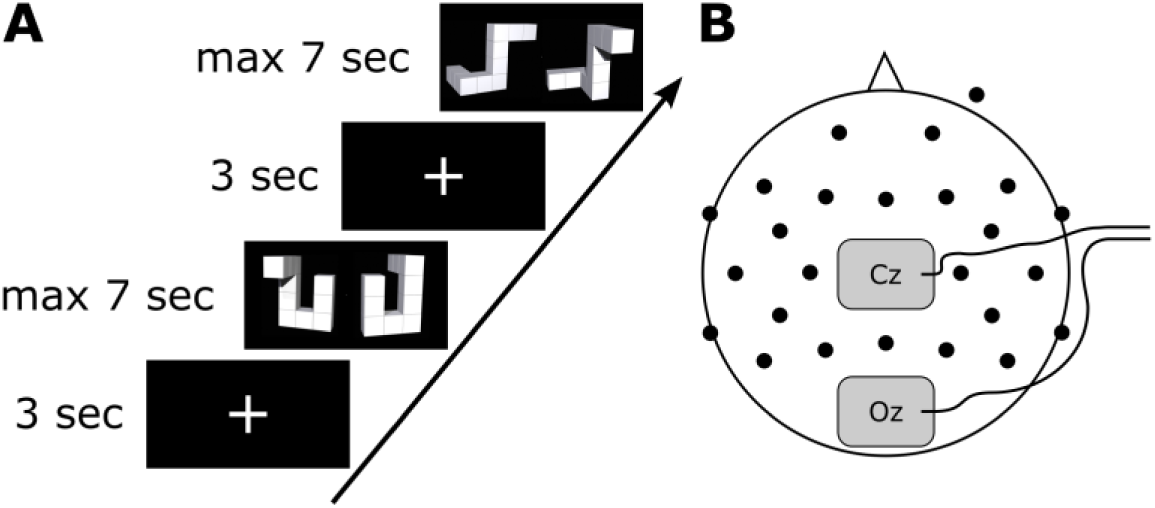
Setup of the human experiment. A: Experimental paradigm showing one trial of the mental rotation experiment. B: Setup of the EEG and stimulation electrodes. Black dots depict EEG electrode positions, grey rectangles stimulation electrode positions.

### 4.1. Materials & Methods

#### 4.1.1. Paradigm

In an electrically shielded, dimly lit room, the subject (male, 26 years, right-handed) was seated in a recliner in front of a computer screen. EEG was recorded during three blocks of the mental rotation task. The task was to judge whether the two presented stimuli are identical although rotated with respect to each other or whether they are different (Ganis and Kievit 2015; Kasten et al. 2018). Each block consisted of 56 stimuli presented for maximally 7 s each, with an inter-stimulus interval of 3 s. Overall, one block took approximately 9 minutes. The three blocks were recorded without tACS, with weak tACS (0.5 mA) and strong tACS (1 mA).

#### 4.1.2. Technical requirements for removing the tACS-artifact from EEG

A most important requirement during EEG recordings is that the amplifiers do not saturate due to the high amplitudes of the tACS-artifact. If that requirement is not met, no artifact correction is possible. The main feature in this regard is the dynamic range of the amplifier: With a 16-bit EEG amplifier, 2^16^ = 65536 amplitude values can be digitized. If every amplitude step represents 0.1 µV, as in the phantom experiment, this results in an amplitude range of ±3276 µV. In that case, the amplitude of the artifact can easily exceed the dynamic range, especially at higher impedances. In our human experiment, this value is already exceeded in some EEG-electrodes at 500 µA stimulator output (see table 3).

While this was not a problem for the phantom, it is desirable to use EEG amplifiers with a wider dynamic range for human tACS–EEG experiments, e.g., 24-bit amplifiers, which would allow for an amplitude resolution of 0.05 µV and a dynamic range of ±419430 µV. We therefore opted for a 24-bit system for the human experiment.

A second factor that has to be taken into account to avoid amplifier saturation is bridging of the tACS electrodes with the recording electrodes. One common technique to reduce the impedance of the tACS electrodes to the scalp is the use of saline-soaked sponge pockets that enclose the tACS electrodes. This bears the danger of leaking saline solution that results in a connection of tACS and EEG electrodes and also of EEG electrodes with each other. To prevent this, we recommend using adhesive paste (e.g., Ten20, D.O. Weaver, Aurora, CO, USA), which does not leak and also prevents electrode movements.

Third, it is of great importance that EEG and tACS are synchronized. Different EEG-systems allow for digital synchronization of the recorder with a different system, e.g. using the BrainVision Syncbox (Brain Products, Munich, Germany). The tACS can by synchronized with the EEG recording by generating the tACS signal digitally and passing that signal through a digital-to-analog converter (DAQ) into the stimulator as we have done in the phantom experiment. Additionally, the stimulation frequency has to be chosen such that the template estimation can be successful: We used 10 Hz, which has a period duration (100 ms) which is an integer multiple of the period of the EEG sampling interval (i.e. 0.1 ms at 10 kHz sampling frequency). At 11 Hz, the cycle length is 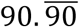ms. Thus at 11 Hz, the zero crossings of the artifact cycles do not coincide with one sample. When estimating a template of the tACS artifact based on one cycle, this leads to an unpredictable error in the template, which in turn leads to a failure of the artifact correction.

#### 4.1.3. EEG

Measurements were performed with a 24-bit battery-powered amplifier (ActiChamp, Brain Products, Munich, Germany) and 24 preamplifier-equipped electrodes mounted in an elastic cap (Acticap, Falk Minow, Munich, Germany) positioned according to the International 10–20 system (Fig. 6B). Electrode impedances were kept below 10 kΩ. The EEG was measured against a common reference at position FP1 and sampled at 10 kHz as in experiment 1. The EEG recording was synchronized with the tACS to guarantee an accurate measurement of the tACS artifact. With the ActiChamp system, it is possible to synchronize the two systems conveniently without a SyncBox.

#### 4.1.4. tACS

The tACS current (10 Hz, with an intensity of either 0.5 or 1 mA), synchronized with the EEG, was applied using a battery-powered stimulator system (DC stimulator plus, Eldith, Neuroconn, Ilmenau, Germany) positioned next to the subject inside the cabin. EEG recording and tACS were both sampled at 10 kHz. Two rubber electrodes (5 × 7 cm), centered at Oz and Cz (corresponding to the stimulation sites of Experiment 1), were attached to the subject’s head using adhesive conductive paste (Ten20, Weaver and Company, Aurora, CO, USA). The tACS signal was created digitally in Matlab and transformed into an analog signal by a NI-DAQ before it was fed into the stimulator as in experiment 1. The stimulator then uses a gain of 2 on its external input to forward the external signal to the subjects’ head.

#### 4.1.5. Correction of the tACS artifact

Since phantom data suggested that SSP would be effective in reducing the artifacts, we expected SSP to correct the tACS artifact also in the human EEG. With a few modifications, the method was directly transferred to the human data. The most relevant difference was that we recorded not only 60, but 600 s of tACS–EEG data, which represents a more realistic scenario in an EEG-experiment. The SSP method relies on an accurate estimate of the template of the artifact. The accuracy of the template, however, depends on the length of the data taken into account: if more repetitions of the artifact (in our case, cycles of tACS) are averaged for the template, more residual EEG activity in the template is averaged out. It is known that the tACS artifact changes in amplitude over time due to changes in impedances between skin and stimulation as well as EEG electrodes. We therefore decided to apply the SSP procedure on portions of EEG data of 15 seconds each, while in the phantom data the entire recording of 60 s was corrected at once. After correction, the data were concatenated such that all analysis procedures could be performed as on the raw data. Other parameters, such as for SIR were identical to those used on the phantom data.

#### 4.1.6. Analysis of human EEG

As expected, the tACS artifact covered brain activity recorded during weak and strong tACS (Fig. 7) with sharp peaks at the tACS frequency, the amplitude of strong tACS artifact being about 2 times higher than weak tACS (Fig. 7B,C). The strong impact of the tACS was also visible in the topographies: While the topography of the average FFT amplitudes at 10 Hz of the EEG without tACS showed an occipital maximum, the topographies of the EEG with tACS represented only the centralized tACS artifact (Fig. 7C). In order to correct the EEG for the tACS artifact, we applied SSP in all conditions, including tACS-free baseline measurements (Fig. 8A,B). Comparing artifact-free EEG before and after SSP indicates a slight amplitude reduction regardless of frequency (Fig. 8). The signal amplitude is attenuated to a similar extent at all frequencies because SSP is a spatial filter. Mutanen et al. (2016) showed that SIR can well restore the original neuronally generated topographies but cannot perfectly compensate for the SSP-introduced attenuations in the signal amplitude. Channels with higher pre-SSP amplitudes also showed a stronger absolute amplitude reduction after SSP (Fig. 8, right).

**Figure 7.**
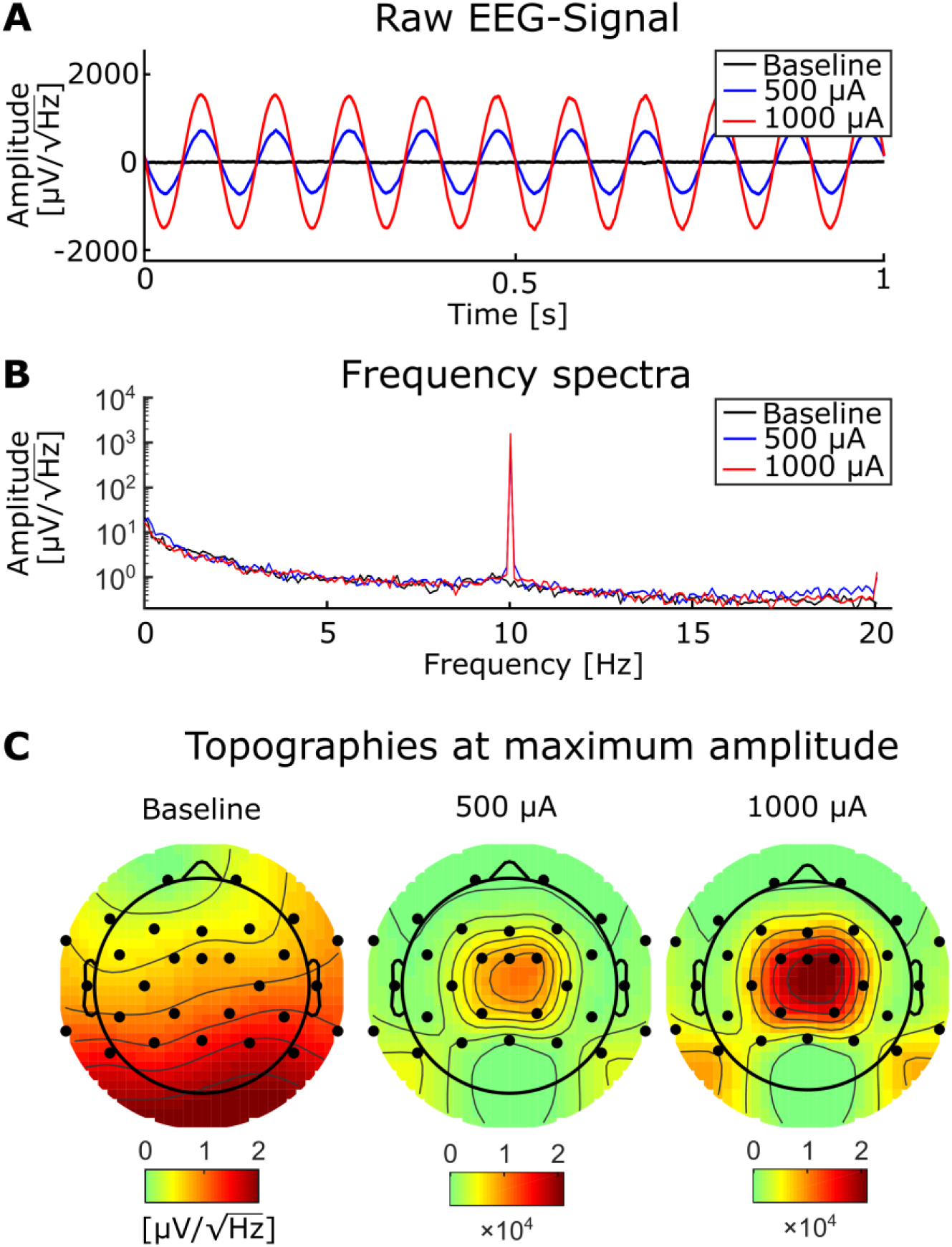
Baseline EEG and tACS-contaminated EEG. A: A representative segment of EEG without tACS (black), and with simultaneous tACS at 500 (blue) and 1000 µA (red). B: Frequency spectra at 0.2-Hz resolution from the same conditions as in A and with the same color conventions. C: Topographies of the baseline EEG (left) and the EEG contaminated by tACS with 500 (mid) and 1000 µA (right).

**Figure 8:**
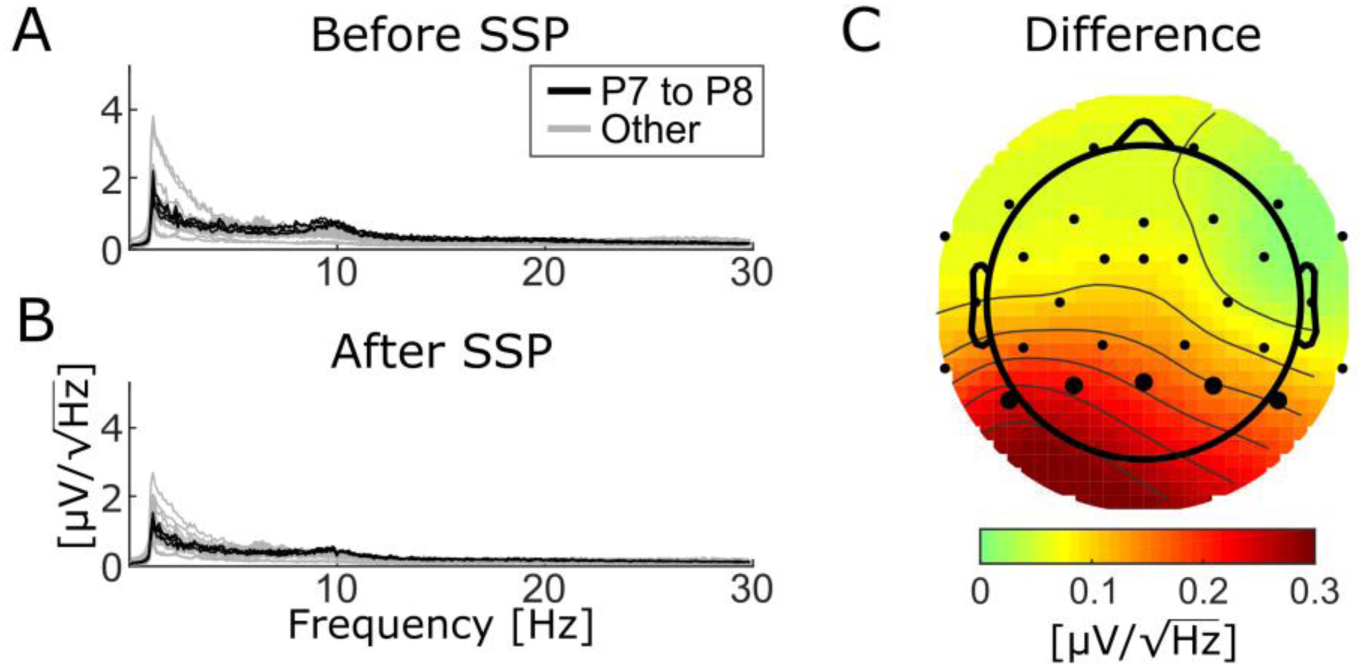
Amplitude attenuation after SSP. Comparison of the tACS-free EEG before (A) and after (B) the SSP algorithm was applied. The alpha peak at 10 Hz is distinguishable before and after but reduced after SSP. Keep in mind that these data represent EEG from a mental rotation task, during which alpha is generally expected to be low. The topography (C) shows the difference in the 8–12-Hz range from before to after SSP correction. Note that there are no occipital channels (O2/O1), and therefore difference values at occipital positions are extrapolated from neighboring electrodes.

After the EEG data were subjected to the SSP algorithm, they were segmented into epochs of 8 seconds around the onset of a mental rotation stimulus (−4 to +4 s around the stimulus). Time– frequency spectra were calculated of each epoch, using wavelets with 7 cycles over the whole time– frequency range. To show the effectiveness of the SSP algorithm, no baseline correction was applied in the frequency dimension, i.e., the TF data were not normalized to a pre-stimulus baseline period. Thus, pre-stimulus activity is visible in the spectra. Furthermore, event-related (de-)synchronization (ERS/ERD) was calculated for the alpha band (8–12 Hz), by computing the absolute difference between pre- and post-stimulus interval as suggested by Kasten et al. (2018):

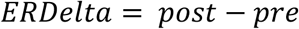

Here, ‘pre’ are ‘post’ correspond to averaged amplitudes between 200 and 50 ms before stimulus onset and between 100 and 500 ms after stimulus onset, respectively. *ERDelta* is positive when the amplitude is increased after the presentation of an (visual) event (ERS). Negative values represent a decrease of the amplitude, i.e., an event-related desynchronization (ERD). This simple subtraction method is preferable to the more established method by Pfurtscheller and Lopes da Silva (1999) that is based on relative change in oscillatory power. When dealing with tACS-contaminated data, relative change can be strongly biased by residual artifacts in the data, while absolute differences are more robust to such influence. Under the assumption that the strength of the tACS artifact is not systematically modulated by the task, residual artifacts after correction can cancel out (Kasten et al. 2018; Kasten and Herrmann 2019).

### 4.2. Results

In the time–frequency (TF) spectra before artifact correction (Fig. 10A), one can see the strong tACS artifact in the second (500 µA tACS) and third row (1000 µA tACS) as a relatively broad red bar which appears unmodulated throughout the time period depicted. Due to its high amplitude, the artifact dominates the TF spectrum. In fact, no characteristics of original brain activity can be seen in these plots. Another feature of these plots is the exaggerated 50-Hz line noise artifact. The enormous strength of which can be explained by the experimental setup: The stimulation signal is transmitted from the DAQ to the stimulator through a BNC cable. Even though those cables are shielded, they capture the line noise via electromagnetic induction. Since the stimulator directly transfers the incoming signal to the stimulation electrodes with a gain of 2, the induced line noise is amplified when the signal is conducted to the human scalp. In turn, the 50-Hz noise is amplified in the EEG-recordings.

In the baseline tACS condition, the alpha decrease (ERD) after stimulus presentation can be seen by visual inspection. For a better comparison, the color bars for all TF spectra depict the same value range. After artifact correction, time–frequency spectra of the tACS conditions (50-µA tACS, 500-µA tACS) did not show an apparent residual artifact (Fig. 9C) and natural alpha fluctuations became visible. Data from the baseline tACS condition has also been subjected to the SSP algorithm. Therefore, the difference between the two time–frequency spectra in the first rows of Fig, 10 A and C show the amplitude reduction due to SSP as described above (Fig. 8). The topographies show ERD over parietal areas before artifact correction (Fig. 9B) and afterwards (Fig. 9D). Note that no occipital electrodes were measured due to the stimulation electrode that covered that area. Thus, shading over occipital areas is extrapolated from other channels. From visually inspecting the topographies, one can clearly identify ERD over parietal areas in all conditions after SSP. We correlated the topographies in all possible combinations and revealed highly significant correlations for all possible comparisons of tACS conditions, i.e., Baseline, 500 µA, and 1000 µA (Table 1). Since these comparisons are not solely between conditions, but also resemble different time points of measurement, the correlations between the corrected data are hard to compare.

**Figure 9:**
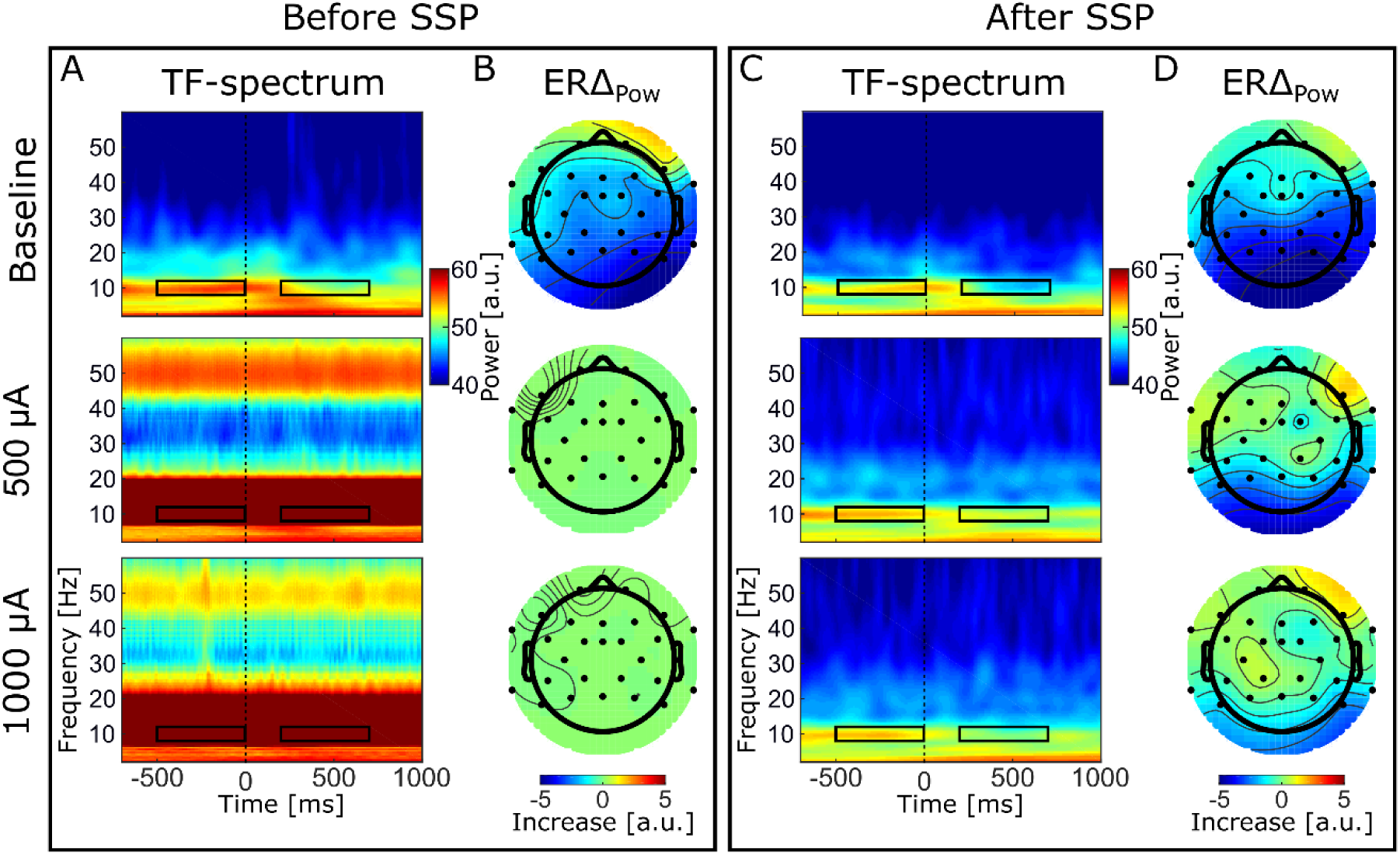
Time–frequency results. Left: before SSP; Right; after SSP. TF spectra depict decompositions of event-related EEG data from electrode Pz. Color bars are the same for all TF spectra (A and C). Black rectangles illustrate the time–frequency sub-spectra used to calculate the ERS topographies.

To provide a comparison of the correlation strengths between conditions after correction, we calculated correlations between topographies from odd and even numbered trials (odd/even split). These values are comparable in size with correlation values between conditions (Table 1). Additionally, we provide the correlation between baseline topographies before versus after SSP which also lies within the same range (*r* = 0.85).

The similarity between the topographies after SSP demonstrate that SSP can successfully recover subtle changes in alpha power on minor scales.

**Table 1:** Correlations between topographies in different tACS conditions and correlations between odd/even splits within tACS conditions as a control. *p*-values were Bonferroni-corrected for multiple comparisons.

| Experimental Conditions | | $\rho$ | $p$ |
| --- | --- | --- | --- |
| Baseline | 500 μA | 0.73 | < .001 |
| Baseline | 1000 μA | 0.53 | .009 |
| 500 μA | 1000 μA | 0.70 | < .001 |
| Control Correlations |  |  |  |
| Baseline odd | Baseline even | 0.77 | < .001 |
| 500 μA odd | 500 μA even | 0.45 | .001 |
| 1000 μA odd | 1000 μA even | 0.68 | < .001 |
| Original Baseline | SSP Baseline | 0.85 | < .001 |

Since ERDelta is a relative measure, one could argue that the correction of the artifact is not even necessary because the calculation of ERDelta is a normalization to a pre-stimulus baseline in the time dimension and would thus be sufficient to cancel out the artifact. However, a recent simulation indicates that strong attenuation of the tACS artifact is necessary to allow the cancellation by computing difference measures to work (Kasten & Herrmann, 2019). To test this assumption on real data, we calculated ERDelta also for the uncorrected data. The resulting topographies are depicted in Fig. 9B. The range of the ERDelta is strongly reduced in comparison to the data after SSP correction (Fig. 9D) in the tACS conditions in rows 2 and 3. Also the topographies do not show the expected occipito–parietal orientation but appear distorted without a distinctive pattern (Fig. 9B, row 2 and 3). In line with predictions of the simulation in Kasten & Herrmann (2019) this result indicates that the artifact completely masks the stimulus-related amplitude reduction and that a reduction of the artifact is necessary to recover the underlying brain activity.

## 5. Discussion

We evaluated artifact-correction techniques to find a feasible method to remove the tACS artifact from EEG data. By comparing different methods applied to phantom data, we found SSP to perform best in recovering the signal of interest. As a proof of concept, we applied SSP to human data from a tACS–EEG experiment and demonstrated to which extent oscillatory parameters such as event-related oscillations can be recovered.

Our initial question was how to estimate that an artifact-correction method does not remove the brain responses of interest and minimizes residual artifacts remaining after correction. We approached this question by applying different methods on a phantom head in which we were in full control over the to-be-recovered EEG signal and the stimulation signal. We found that the SSP method performed best with only minor distortions of the EEG signal compared to template subtraction and sine fitting. For the phantom head, we could quantify this distortion, finding that SSP mildly overcorrects the artifact. However, with SSP this overcorrection can be taken into account when comparing the cleaned signal to the baseline (See supplementary material for details.). With the phantom, possible physiological effects might be underestimated, because the SSP correction attenuates endogenous amplitudes at the stimulation frequency. For human data, we cannot be entirely sure about the performance of the correction as we do not exactly know the ground truth; however, we compared the same experimental conditions with and without concurrent tACS. Overall, these results suggest that the artifact correction was successful despite an overall reduction of amplitudes; SSP was able to recover subtle changes in alpha amplitude (ERD) relative to a pre-stimulus baseline.

### 5.1. Phantom vs. human head

An important question is whether the results obtained in the phantom experiment can be generalized to human data. Obviously, a human head is not three-layered and perfectly spherical. A study by Kim et al. (2015), however, showed that the three-layer spherical model is quite accurate in capturing the essential characteristics of the electric-stimulation-generated ohmic currents in scalp, skull and the brain. Another difference lies in the sources of activity: Contrary to the many sources in a human brain, our phantom only had one neural source. Given that SSP performance depends on the orientation and location of the neural source, the method might further attenuate some neural signals of interest given that their topography can be similar to that of the artifact. This problem can be tackled by evaluating possible signal distortions after SSP (Uusitalo and Ilmoniemi, 1997).

In our phantom study, the dipolar current source oscillated independently of the external stimulation. This might produce over-optimistic results when using template subtraction. If neural activity synchronizes to the tACS, it will be attenuated after subtracting the template; however, perfect phase alignment cannot be expected from real neuronal activity (van Veen et al., 1997). This also applies to the sine-fitting method: only if the tACS entrains perfectly to the neural frequency at which the brain is stimulated, then sine fitting will also attenuate the entrained brain oscillations.

Another difference of the phantom measurement compared to real human EEG data concerns the impedance: In the phantom head, the artifact amplitude was constant over the course of the stimulation because the electrode–skin impedances remained constant. This allowed us to use all trials to create a template to be subtracted. For real human EEG data, this is not necessarily the ideal approach because the tACS artifact amplitude varies over time due to changes in impedance, elicited by physiological processes in the human body such as heart-beat and respiration (Noury et al., 2016). This poses a problem for the template subtraction method because the subtraction of an incorrectly sized template will result in a residual artifact: the fewer trials are used to create the template, the wider the notch in the Fourier spectrum will be. The problem notably also applies to the SSP method, since the artifact subspace is also calculated based on a data-based template of the artifact. In the human experiment, we tackled the problem of varying impedances by applying the SSP in temporal step of 15 sec (see Section 4.1.5).

### 5.2. Sine subtraction

Most commonly, the tACS signal is a sine wave (Herrmann et al., 2013). Therefore, it is an intuitive assumption that one can simply fit and subtract a sine from the contaminated EEG signal and the artifact is removed. An advantage of this method would be that the signal in each electrode can be cleaned separately; this may be beneficial in experimental setups with a small number of electrodes; however, we demonstrated that the sine subtraction method shows a comparatively poor performance. The main problem with sine fitting is that using a least-squares criterion can result in overfitting or underfitting if a significant proportion of the EEG signal of interest phase aligned to the artifact. Another problem with sine fitting is that the tACS artifact is not a perfect sinusoidal wave, but rather a series of analog amplitudes generated by a digital-to-analog converter (DAC), i.e., the sine wave is approximated by a kind of step function and each step is superposed with an exponential due to capacitance inside the DAC. Additionally, the measured artifact is non-sinusoidal due to its interaction with physiological tissues (Noury et al., 2016). A perfect sine wave subtracted from a slightly distorted sine wave will result in a residual artifact. If the artifact is several orders of magnitude larger than the neuronal signals of interest, even small relative differences between the perfect sine-wave model and the actual tACS artifact waveform can cause large absolute errors in the corrected EEG. Overall, our results suggest that this subtraction method cannot be recommended for tACS artifact correction.

### 5.3. Template subtraction

Like the sine-wave-subtraction method, template subtraction can remove the artifact for each electrode separately. Compared to the sine-fitting approach, template subtraction demonstrated a clearly better performance at recovering the baseline signal in the phantom data; however, especially the temporal fine structure (phase) could not be perfectly recovered. Before applying template subtraction to human tACS–EEG data, several practical considerations should be taken into account.

Typically, the size of the tACS artifact in human EEG can vary due to impedance changes of the tissue (Noury et al., 2016), which can result in improper templates and subsequent residual tACS artifact or a loss of neural EEG signal. Fitting the template to the artifact in the raw EEG by minimizing the sum of squares (Helfrich et al., 2014) can help with the problem of variation in the amplitude of the artifact; however, this can also result in over- or underfitting. In the phantom-head data, we found that fitting the template to the artifact using least squares resulted in worse recovery of the contaminated signal than simple template subtraction (data not shown). Another solution would be to use temporarily more specific templates by averaging a smaller number of adjacent cycles (moving-average approach); however, the less cycles are included to compute the template, the wider the affected frequency range.

### 5.4. SSP

We found SSP to yield the best artifact-correction performance. Artifact-contaminated phantom data could be recovered almost perfectly; the application to human data is promising. A major difference between SSP and both sine fitting and template subtraction is that it is based on spatial filtering, thus it may project out artifactual components that are invisible to sine fitting and template subtraction. Even though SSP is able to correct the artifact almost completely in the phantom data, it distorts signals although in a perfectly known way: the cleaned signals are not meant to be estimates of the signal in the original channels in question (Mäki & Ilmoniemi, 2011). The signals after SSP are known linear combinations of the original EEG signals and can be used without bias in source estimation (Uusitalo & Ilmoniemi, 1997) as long as the data still has a sufficient dimensionality. SIR can correct some of the SSP-induced spatial distortions; however, the original signal amplitudes cannot be perfectly recovered in all channels because some linear components of the signals have been zeroed, leading to overall reduction in amplitude (Fig. 8). We were able to recover time–frequency spectra showing ERD in the alpha range after correction. This indicates that SSP does not completely diminish activity at the stimulation frequency like a notch filter would, but can recover activity even at the stimulation frequency. Comparing tACS-free data with and without application of SSP–SIR reveals a general decrease of amplitudes in the FFT-spectra, which seems to be stronger at higher amplitudes. Topographic similarity of the artifact and the signals of interest contribute to the unwanted attenuation of the latter; the more similar they are, the higher the attenuation. This is also evident in the human data as the topography of the resting-state alpha and the artifact topographies varied significantly. We advise to apply the SSP–SIR method when comparing data from tACS conditions with tACS-free conditions. Still, most reliable results can be achieved when contrasting experimental conditions combined with the same tACS condition. There is some consensus among different research groups that under the assumption that residual artifacts are present in two experimental conditions to a similar degree, they can cancel out when computing difference measures, such that only the physiological effects remain (Neuling et al., 2015; Kasten et al. 2018, Noury & Siegel 2018, Herring et al. 2019). In line with predictions of a previous simulation (Kasten & Herrmann, 2019), our data demonstrate that it is insufficient to contrast uncorrected data: although the artifact is constant over time, a baseline correction in the form of a subtraction in the time-frequency space, could not reveal the ERD while the SSP method was able to recover these subtle changes in alpha amplitude.

### 5.5. General

Overall, our results in human data indicate that the SSP method attenuated the tACS artifact sufficiently strong to recover task related modulations of endogenous brain oscillations. This is supported by the observation that although the artifact covers brain activity in all EEG channels before correction, after SSP the task induced alpha power modulation is strongest in parietal channels, resembling the topography of the artifact free data. It should be noted, however, that this does not imply that the tACS artifact has been removed entirely from the human EEG recordings. Previous studies have shown, that a variety of physiological processes can give rise to non-linear modulations of the tACS artifact, which can hinder complete tACS artifact removal (Noury et al. 2016; Noury and Siegel 2017). Future studies will need to evaluate to which degree these non-linearities affect tACS artifact cleaning performance of the SSP method. Nevertheless, our results already indicate that SSP might be a powerful alternative to template subtraction for the analysis of concurrent tACS-EEG data, as the latter suffers from overcorrection (Helfrich et al. 2014) and insufficiently accounts for non-linear modulations of the tACS artifact (Noury et al. 2016).

The SSP method should be further explored in future studies to find the best template the artifact subspace is estimated on. In the current study, the SSP operator was computed from the average template, which might contain entrained brain signals (Thut et al., 2011). As a consequence, this brain activity would also be removed. This issue could be overcome by estimating the artifact subspace based on different tACS conditions (e.g., two or more frequencies and amplitudes), thus minimizing the contribution of brain activity to the template and maximizing the contribution of the artifact.

## 6. Conclusion

SSP yielded by far the best performance in removing the tACS artifact and recovering the brain activity in EEG recordings in comparison to template and sine-wave subtraction. Even though the performance on the phantom cannot be unequivocally extended to human EEG measurements, SSP is a strong candidate for the correction of the tACS artifact in combined tACS–EEG studies.

## 7. Acknowledgements

The study was supported by the German Research Foundation (Deutsche Forschungsgemeinschaft, DFG), grants SFB/TRR 31 and SPP 1665 (CSH), Academy of Finland, and the Finnish Cultural Foundation.

## 9. Author Contributions

TN, TPM, JV, RJI, CSH designed the experiments. TN performed the phantom experiment, analyzed the data, and wrote the paper. TPM built the phantom head, analyzed the data, and wrote the paper. JV performed the human experiment, analyzed the data, and wrote the paper. RJI and CSH wrote the paper.

