## Supplemental Materials for "Signal-space projection suppresses the tACS artifact in EEG recordings"

### 11. Supplemental Material and Methods

#### 11.1. Phantom head construction

The phantom head consisted of two spherical polycarbonate shells with diameters of 120 and 140 mm, with a shell thickness of 2 mm. Similar material was used by Höfner et al. (2011) when studying the ohmic currents produced by a dipolar current source within a spherical object. Both shells consisted of two equally sized hemispheres, which we fixed together with cyanoacrylate (Loctite Precision Super Glue, Henkel AG & Co.KGaA, Düsseldorf, Germany). The two shells were then attached to each other to form a concentric, three-layer structure (Fig. 1).

The smaller sphere was supported from the bottom by four nylon screws that were tightened to holes drilled to the bottom of the larger shell. The largest screw (6 mm wide) was fastened to the south pole of the sphere and the three smaller ones (3 mm wide) were equally spaced being 30 mm away from the South Pole. The smaller sphere was glued to the screws with the cyanoacrylate. The same glue was also used to strengthen the attachment between the screws and the outer spherical shell.

We used 1-mm-thick and 25-mm-long copper-wire segments as electrodes. We attached 22 electrodes on approximate positions of the 10–20 EEG system, on the upper hemisphere of the outer spherical shell so that a 1-mm segment of each electrode wire protruded into the outer sphere (see Fig. 1D). The electrodes were inserted and glued to 2-mm-wide holes with a two-component epoxy glue (Plus Schnellfest, UHU GmbH & CO. KG, Bühl, Germany).

The phantom had a dipolar current element to represent a neuronal current source. The dipole was constructed out of paired cable endings (0.2-mm-thick enameled copper wire) that protruded into the inner shell via two 1-mm holes. The dipolar current source was created by opening a 10-mm-long segment of the paired cable and scratching off the insulating coating from the tip of both wire endings (approx. 1-mm segments). The structure was toughened and fastened to the holes with the epoxy glue.

The dipole was placed on the left hemisphere, on the line between P3 and T5, 10 mm from P3 towards T5, inside the skull (10 mm from the outer surface of the "brain"). Both ends of the dipole were tangential with respect to the skull so that they had an anterior–posterior orientation. After the structure of the phantom was finished, all the seams were sealed with waterproof glue (Casco LiquiSole, Sika Finland AB, Espoo, Finland).

To mimic the conductive properties of the human skull, we drilled 2000 holes with diameter of 1 mm through the inner shell. Via this procedure, the inner shell had a conductivity that is roughly 1/30th of the outer sphere. The holes were randomly and approximately evenly distributed over the spherical surface. Finally, the whole concentric structure was filled with NaCl solution through a 4-mm wide hole on the top of the phantom head using a syringe with a 1.7-mm-thick metal needle. After filling the phantom head, this hole was sealed with a rubber cap. The NaCl solution consisted of 3 g of NaCl mixed with one liter of demineralized water resulting in a conductivity of 0.57 S/m in the brain and the scalp.

Because the current tends to travel the shortest path through the insulating skull and because the well-conducting scalp spreads current components coming through each hole widely across the scalp, we can approximate that for the purpose of EEG the constructed skull appears to have a constant conductivity. This conductivity is determined by the ratio between the total area of the holes and the surface area of the smaller spherical shell. Thus the conductivities of the three layers were 0.57 S/m, 0.019 S/m, and 0.57 S/m for the “brain,” “skull”, and “scalp”. The true conductivities of these tissues are still not well known, but the values here fall well within the variation of most common tissues (Gabriel et al., 2009). The ratio between conductivity of the skull and the brain, which is more important in EEG than the absolute conductivity values, corresponded, e.g., with the findings of Gonçalve et al. (2003) and Lai et al. (2005).

11.2. SSP details

Let us consider an EEG dataset $Y_{tACS}$ that has been measured during tACS. The measured

EEG dataset consist of the following components:

$Y_{tACS} = GS_{tACS} + A + N$, (SE1)

where $S_{tACS}$, A, N represent the brain activity during stimulation, tACS-artifacts, and noise signals, respectively. G is the so-called gain or lead-field matrix whose term $G_{ij}$ describes the sensitivity of EEG channel i to neural source j.

If we can assume that the noise and brain signals do not average well across the tACS periods, we can estimate the tACS-artifact topographies from the averaged EEG and project out these topographies using SSP. We write the averaged data in terms of singular-value decomposition:

$Y_{ave} = USV^T$, (SE2)

where the column vectors of U, $u_k$, give all the spatial patterns needed to fully explain the averaged data, and S is a diagonal matrix whose diagonal terms $\sigma_k$ are the singular values that describe how much of the total data variance of the averaged data is explained by each spatial pattern $u_k$. The column vectors of V are the right singular vectors, which are not needed for SSP.

We suppress the tACS artifact by projecting out the K most prominent (based on their data variance) spatial patterns of $Y_{avev}$ from $Y_{tACS}$ with an operator

$$P = I - [u_1, u_2, \dots, u_k][u_1, u_2, \dots, u_k]^T, \text{ (SE3)}$$

which approximately fulfills

$$PA \approx 0. \text{ (SE4)}$$

This operation removes all the information that can be explained with the spatial patterns $u_k$, k=1,2,…,K. If the spatial pattern of the artifact does not change over time, K=1 is sufficient for removing the tACS artifact. Otherwise, more components need to be removed. After SSP, the data become:

$$PY_{tACS} = PGS_{tACS} + PA + PN = PGS_{tACS} + PN \quad \text{(SE5).}$$

Notice, that also the brain and noise signals will be modified to some extent. This can be taken into account with source-informed reconstruction (SIR) (Mutanen et al. 2016) by multiplying the data with a reconstruction operator R. Thus, the cleaned data can be written:

$$Y_{clean} = RPY_{tACS} = RPGS_{tACS} + RPN, \text{ (SE6)}$$

where $R = G(PG)^{\dagger}$, $(\blacksquare)^{\dagger}$ standing for the pseudoinverse.

Even after SIR, SSP attenuates the signals in different channels to some extent. However, analyzing entrainment is still possible because we are usually more interested in the relative changes of oscillatory activity compared to baseline. Since we know exactly the linear operation involved in the SSP cleaning, we can take this information into account when comparing the tACS–EEG signal to the baseline EEG.

Let us consider that we have also performed a baseline measurement that quantifies spontaneous oscillatory activity. The baseline data $Y_{base}$ can be written as:

$$Y_{base} = GS_{base} + N, \text{ (SE7)}$$

where $S_{base}$ contains neural activity during the baseline measurement.

If we now examine the expressions for cleaned and baseline data (Eqs. (SE6) and (SE7)), we see that to demonstrate entrainment, we cannot directly compare the cleaned tACS data to the baseline data. This is because instead of the original gain matrix G, we have the RPG term describing the sensitivity of each cleaned EEG channel to different neural sources.

To compensate for the distortion introduced by the transform RP, we can apply the same transformation to our baseline data:

$$\mathrm{RPY}_{\mathrm{base}} = \mathrm{RPGS}_{\mathrm{base}} + \mathrm{RPN}. \qquad \text{(SE8)}$$

If we now observe a change from $\mathrm{RPY}_{\mathrm{base}}$ to $\mathrm{Y}_{\mathrm{clean}}$, we know it is likely to result from a difference in brain activity. This is an advantage of SSP because it allows to take into account the possible overcorrection when comparing the cleaned and the baseline data.

### 11.3. References (Supplemental)

Gonçalve, S., De Munck, J. C., Verbunt, J., Bijma, F., Heethaar, R. M., & Lopes da Silva, F. (2003). In vivo measurement of the brain and skull resistivities using an EIT-based method and realistic models for the head. *IEEE Transactions on Biomedical Engineering*, 50(6), 754–767.

Gabriel, C., Peyman, A., & Grant, E. H. (2009). Electrical conductivity of tissue at frequencies below 1 MHz. Physics in Medicine and Biology, 54(16), 4863.

Höfner, N., Albrecht, H.H., Cassará, A.M., Curio, G., Hartwig, S., Haueisen, J., Hilschenz, I., Körber, R., Martens, S., Scheer, H.J., Voigt, J., Trahms, L., & Burghoff, M. (2011). Are brain currents detectable by means of low-field NMR? A phantom study. *Magnetic Resonance Imaging*, 29(10), 1365–1373.
